# Leaf specific overexpression of a mitochondrially-targeted glutamine synthetase in tomato increased assimilate export resulting in earlier fruiting and elevated yield

**DOI:** 10.1101/2022.06.28.497938

**Authors:** José G. Vallarino, Sanu Shameer, Youjun Zhang, R. George Ratcliffe, Lee J. Sweetlove, Alisdair R. Fernie

## Abstract

To identify potential strategies for increasing the efficiency of tomato leaf metabolism, with a focus on the links between nitrogen/carbon metabolism, we explored a diel Flux Balance Analysis (FBA) model of a source leaf in which the metabolic output was varied up to the theoretically-achievable maximum. We noticed a potentially interesting switch in the use of glutamine synthetase (GS) isoforms –from the chloroplast isoform to the mitochondrial one- for nitrogen assimilation. To further explore this prediction, we characterized transgenic tomato plants over-expressing two tomato *GS* genes, *GS1* and *GS2*, targeted to mitochondria. Both sets of transgenic plants were characterized as displaying faster growth rate, early flowering and increased fruit yield. In leaves, metabolomic profiling and enzyme activity analysis pointed that GS activity in mitochondrial plays a role in increasing the intracellular synthesis and subsequent export of sugar. Consistent with these changes, higher sucrose concentration in leaf exudates and reduced activities of enzymes involved in leaf starch synthesis were observed. Moreover, mitochondrial GS activity affected chloroplast redox status in a manner that modulated photorespiration and nitrogen metabolism. The combined data reveal the influence of mitochondrial GS activity on both foliar carbon/nitrogen balance and regulation of source-sink metabolism in tomato plants.

## INTRODUCTION

Research on fruit development and ripening in crop species that have agricultural value is essential given that these processes influence yield, quality and harvesting efficiency (Klee and Giovannoni, 2011; Seymour et al., 2013; Kwon et al., 2020; Martín-Pizarro et al., 2021; Ren et al., 2021). Similarly, advances have been made recently in understanding the mechanisms that link metabolism to crop yield (Riedelsheimer et al., 2012a; Riedelsheimer et al., 2012b; de Abreu et al., 2017) including fruit yield (Schauer et al., 2006; Schauer et al., 2008; Do et al., 2010). That said, there is considerable need to better understand metabolic factors that affect fruit yield such that they can be targeted to improve breeding strategies or genetic manipulation. Within this context considerable recent evidence has accrued that highlights the importance of nitrogen (N) metabolism with respect to carbon (C) partitioning rendering metabolites at the interface of C/N metabolism as attractive targets for the manipulation of source-sink balance (Santiago and Tegeder, 2016, 2017; Tegeder and Masclaux-Daubresse, 2018; Fernie et al., 2020; Lu et al., 2020). That C and N metabolism are strongly intertwined is known at many levels, for example amino acids are vital to the assembly and operation of the photosynthetic apparatus while photosynthesis and photorespiration provide the energy, reductants and C skeletons for N reduction and amino acid synthesis. C positively regulates N assimilation and metabolism at transcriptional and post- translational levels; for example, the control of nitrate reductase (NR) expression (Vincentz et al., 1993) as well as its activity by phosphorylation and interaction with inhibitor proteins (Bachmann et al., 1996). That said legumes with increased N acquisition, source-to-sink transport of organic N, seed development and NUE, suggest that C availability does not always present a hurdle to such improvements (Zhang et al., 2015; Santiago and Tegeder, 2016), given that plants are able to adjust their C metabolism and transport in response to N availability and sink strength (Zhang et al., 2015; Santiago and Tegeder, 2016). In addition, in *Arabidopsis* or other oilseed plants, enhanced amino acid transport towards source leaves and accompanied improvements in photosynthetic NUE can lead to increased seed C supply and ultimately yield, even under N stress conditions (Perchlik and Tegeder, 2018). Intriguingly, flexible adjustment in C and N metabolism and allocation may also additionally occur in the reverse situation, at least under high N availability. A recent study in pea shows that enhanced sucrose partitioning from source to sink results in modifications in leaf C and N metabolism as well as amino acid transport to sinks in order to accommodate the increased assimilate demand of growing seeds (Lu et al., 2020). Indeed, taking such a combined strategy whereby both C and N metabolism are simultaneously altered appears particularly prudent given difficulties in enhancing tomato fruit yield following multi-gene transformation strategies (Vallarino et al., 2020).

Despite the fact that considerable advances and notable successes have been made in engineering plant metabolism (Zhu et al., 2017; Mangel et al., 2019; Vallarino et al., 2020; Hsieh et al., 2021) our ability to rationally manipulate central metabolism that provides constituents for biomass and other outputs remains limited, largely because of the highly interconnected nature of the central metabolic network. It is therefore essential to approach metabolism from a systematic perspective when developing hypotheses for metabolic engineering (Sweetlove et al., 2017) and computational models are central to this endeavour (Küken and Nikoloski, 2019; Clark et al., 2020). For large networks like central metabolism, flux balance analysis (FBA) has emerged as a robust approach to make accurate predictions of fluxes and as a useful in-silico tool to explore network capabilities and identify engineering strategies (Kruger and Ratcliffe, 2015). A constraint-based modelling approach, is increasingly being used to model metabolism in many plant tissues (Grafahrend-Belau et al., 2013; Colombié et al., 2015; Yuan et al., 2016; diCenzo et al., 2020; Shameer et al., 2020) and in specific cell types (Robaina-Estévez et al., 2017; Tan and Cheung, 2020).

In the current study, we used a diel FBA model (Cheung et al., 2014) of metabolism of a tomato source leaf to explore potential flux distributions in the metabolic network as the system approached its theoretical-maximum in terms of efficiency (in this case the amount of sugars and amino acids that could be exported from the source leaf for a given rate of photosynthetic C assimilation). One of the interesting changes we noted at the intersection of N and C metabolism involved glutamine synthetase (GS), with the model suggesting an advantage in boosting the capacity of mitochondrial GS. In plants, there are two GS isoforms, a cytosolic form (GS1) and a plastidic form (GS2) (Bernard and Habash, 2009) that catalyze the ATP dependent synthesis of Gln from NH_4_^+^ and Glu. Taira et al., (2004) in Arabidopsis has pointed that GS2 form functions in both leaf mitochondria and chloroplast to facilitate NH_4_^+^ recovery during photorespiration. Moreover, the SUBA4 database suggested that there is also proteomic evidence for the presence of GS2 in Arabidopsis mitochondria (Hooper et al., 2017).

In this work, we generated two set of transgenic tomato lines (cv MoneyMaker) exhibiting either increasing expression of *GS1* or *GS2* in mitochondria under the control of leaf- and mesophyll-specific promoter, ribulose-bisphosphate carboxylase (RbcS), which confers leaf -mesophyll specificity, respectively. We carried out an extensive phenotypic and molecular analysis of these lines encompassing analysis of primary metabolite levels as well as, expression profiles and cellular fluxes of these lines. The integrated analysis of these comprehensive set of data allows us to discuss these data with respect to current models of the integration of N and C metabolism during plant growth and fruit ripening and development. This approach thus clearly represents a novel strategy for agriculture as such highlighting the power of combining computational modelling with experimental research within approaches to metabolically engineer organ growth.

## RESULTS

### *In silico* exploration of efficiency of the tomato source leaf metabolic network

In order to identify points at the intersection of N and C metabolism that could be potential targets for improving the overall efficiency of leaf metabolism, we constructed a diel stoichiometric model of central metabolism in a tomato source leaf and predicted fluxes in the network using parsimonious flux balance analysis (pFBA). The model was required to produce sugars and amino acids in specified relative proportions as observed in tomato phloem sap (Walker and Ho, 1977; Valle et al., 1998). Non-growth-associated maintenance costs were also accounted for according to the incident light intensity (Töpfer et al., 2020). Further details of model set-up can be found in the Methods section and a full list of constraints applied to the model is provided (Supplemental Table S1). Our strategy to identify fluxes of interest was to perform a constraint scan, varying the output of the model (export of sugars and amino acids to the phloem) from the maximum predicted by pFBA. To do this, the output reaction flux was reduced from the maximum in 2% steps, each time re-running the model and storing the flux predictions. The sets of flux predictions obtained were plotted against model output to identify reactions that show marked changes in behavior as the model approached the theoretical maximum output. Flux- variability ranges were computed for reactions of interest to account for the variability of fluxes within the model solution space. A full dataset of the flux distributions from this process is provided in Supplemental Data Set 1. Of the reactions involved in N assimilation in the leaf, we noticed a potentially interesting behavior with the glutamine synthetase (GS) reaction (Supplemental Figure 1). The output of the model was positively correlated with overall GS reaction flux, which is not surprising as the output includes amino acids whose synthesis requires the assimilation of nitrate in the leaf. But additionally, there was a marked switch in the choice of GS isoform used as the model reached approximately 90% of the theoretical maximum. Up to that point, N was assimilated exclusively using the chloroplast GS. Beyond that point, the contribution of the chloroplast GS began to decline, and the mitochondrial GS took over. Very close to the theoretical maximum (∼97%) a small flux in cytosolic GS was predicted, although flux variability analysis indicated that this small flux could be entirely taken up by additional mitochondrial GS with no detriment to the output (Supplemental Figure 1). These predictions suggest that there may be an overall efficiency gain by introducing a functional GS into leaf mitochondria.

### Mitochondrial over-expression of *GS1* and *GS2* in tomato (cv. MoneyMaker) leaves

Full-length cDNAs encoding *GS1* (*Solyc04g014510*) and *GS2* (*Solyc01g080280*) were isolated by a PCR-based approach. A 1071 bp fragment of the cDNA encoding GS1 was cloned in the sense orientation into the transformation vector pK2GW7 under the control of the leaf mesophyll- specific promoter (RbcS) and in frame with the mitochondrial transit peptide COX4ts (which results in a mitochondrial localization of the encoded protein) (Nelson et al., 2007). Similarly, a 1316 bp fragment of the cDNA encoding GS2 was cloned in the sense orientation behind the RbcS promoter and the COX4ts mitochondrial transit peptide. Using an established *Agrobacterium tumefaciens*- mediated gene transformation protocol, we were able to generate ten and fourteen independent transgenic tomato lines harboring *RbcS-COX4ts-GS1* and *RbcS-COX4ts-GS2* constructs, respectively. The expression levels of *GS1* and *GS2* were subsequently analyzed in the leaves. Expression was detected in all transgenic lines and the untransformed wild type (control; data not shown). Four lines per construct were chosen: GS1-2, GS1-6, GS1-33, and GS1-3 presenting low, medium, and high overexpression (RbcS-COX4ts-GS1 lines) and GS2-14, GS2-19, GS2-21, GS2-23 (RbcS-COX4ts-GS2 lines) which displayed more homogeneous overexpression between lines (Figure 1A). Analysis of GS activity revealed that these lines displayed a significant activity increases between 1.2- to 1.8-fold in leaves (Figure 1B).

**Figure 1.**
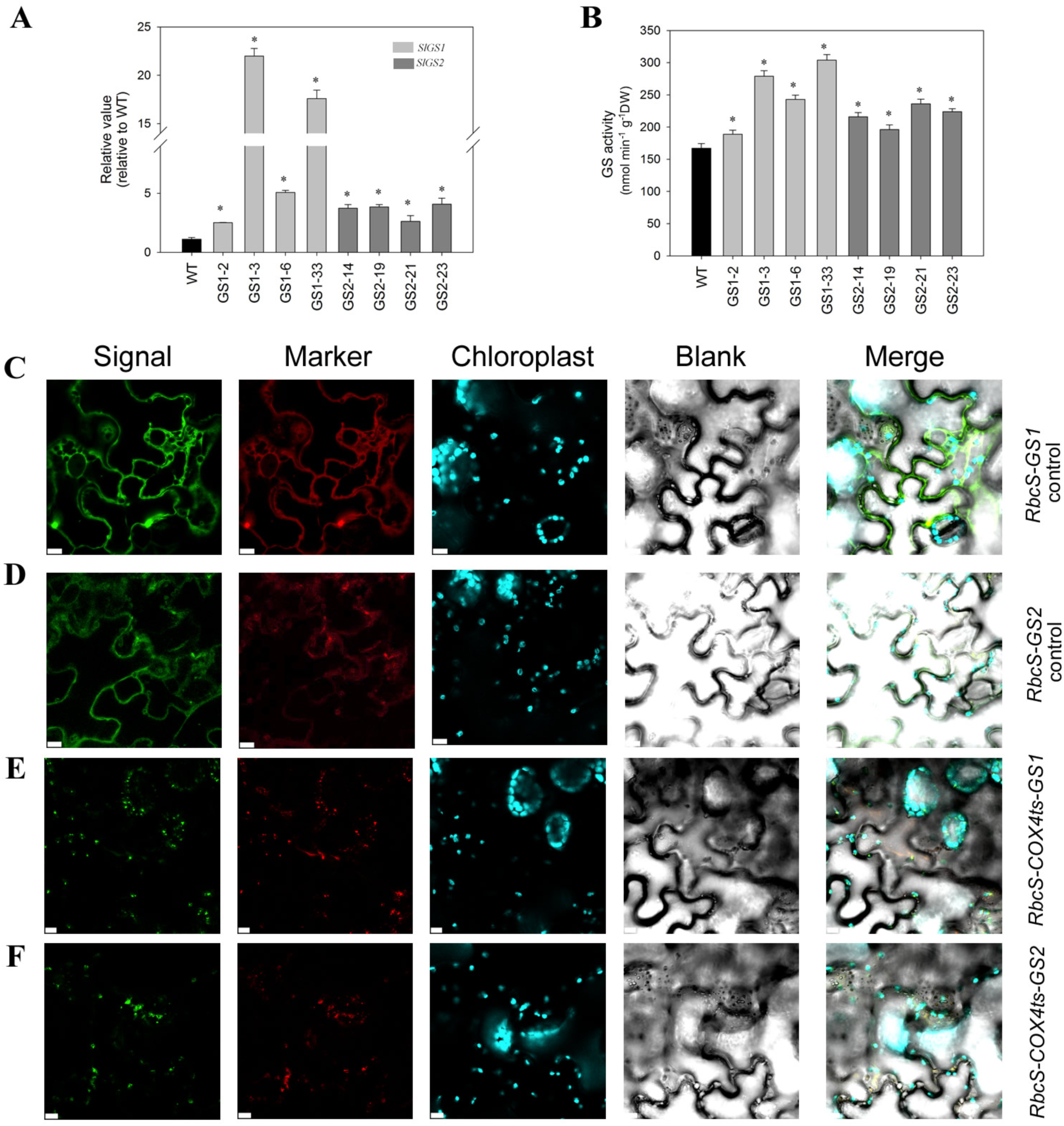
Preliminary characterization of transgenic lines. Subcellular localization of RbcS-COX4ts-GS1 and RbcS-COX4ts-GS1. **(A)** Expression of tomato *SlGS1* and *SlGS2* genes by qRT-PCR in source leaves. **(B)** Total GS activity in source leaves (enzyme activity was normalized to dry weight). For A and B, data were obtained from the third fully expanded leaf from the apex of six-week-old plants. Leaves were harvested at middle day. Values are means from five replicates ± standard error. Statistically significant differences by Student’s *t* test are indicated by asterisk (*P*<0.05). **(C-F)** Mitochondrial targeted *SlGS1* and *SlGS2* were analyzed by transient expression in *N. benthamiana* leaves. The signal of the expression of control constructs RbcS- GS1 **(C)** and RbcS-GS2 **(D)** present the cytosolic expression with cytosolic mCherry marker **(C)** and mCherry plastic marker **(D)**. The signal of the expression of constructs RbcS-COX4ts-GS1 **(E)** and RbcS-COX4ts-GS2 **(F)** present the mitochondria sub-cellular localization with mitochondrial mCherry marker. Merge signals are indicated above the panels. Bar, 10 μm.

Sub-cellular targeting was tested by transient expression experiments in which both coding regions (CDS) of GS1 or GS2 and the CDS fused to the mitochondrial transit peptide COX4ts were linked with fluorescent GFP. The cytosol, plastid, and mitochondria mCherry constructs were used as the marker of these organisms. The transient agro-infiltration method developed by Zhang et al., (2020) was used to infiltrate *Nicotiana benthamiana* leaves and after 3 days, fluorescence was examined by confocal microscopy. As described in previous works, the native GS1 was largely subcellularly localized to the cytosol while GS2 was present in both the cytosol and plastid. However, when the mitochondrial signal peptide was fused to either GS1 or GS2 there localization was only in correspondence with the mitochondrial marker (Figure 1C-F). In order to assess if this was also the case in the transgenic lines, we isolated mitochondria from the wild type and the strongest overexpressing line of both GS1 (line GS1-3) and GS2 (line GS2-14) and determined the activities of GS as well as the marker enzymes UDP-glucose pytophosphorylase, UGPase (cytosol) and ADP-glucose pyrophosphorylase, AGPase (chloroplast) as well as the additional mitochondrial enzyme succinate dehydrogenase, SDH, in order to account for the loss of mitochondria during the isolation process. Despite very little contamination with choroplastic or plastidic material (latencies of either enzyme were under 3% in all cases), the GS activity was significantly enriched in purified mitochondria of the transgenics (being 10.5±3.2, 41.2±2.5 and 29.1±3.1nmol min^-1^ gFW^-1^ for wild type, GS1-3 and GS2-14, respectively). Indeed, although marginally higher in the mitochondria the increase in mitochondrial GS activity was not significantly different to the total cellular increase of GS activity measures in this harvest (compare 30.7 and 18.6 nmol min^-1^ gFW^-1^ as mitochondrial GS activity increases in lines GS1-3 and GS2-14 to cellular GS activity increases of 29.1 and 17.2 nmol min^-1^ gFW^-1^ in the same lines, respectively). These data alongside the GFP localization data thus provide compelling evidence that the transgene product was successfully and completely targeted to the mitochondria.

### Leaf specific over-expression of mitochondrially targeted *SlGS1* or *SlGS2* results in increased plant biomass and fruit yield

In order to assess the effect of mitochondrially targeted *GS1* and *GS2* overexpression across the growth and development periods of tomato plants, a detailed phenotypic analysis was performed. Interestingly, when transgenic plants were growth in greenhouse side-by-side with untransformed controls, we observed a clear difference in growth at the beginning of development. Both sets of transgenic lines, *RbcS-COX4ts-GS1* and *RbcS-COX4ts-GS2,* displayed a significantly faster growth rate, with *RbcS-COX4ts-GS1* lines being slightly taller than *RbcS-COX4ts-GS2* plants, but both being taller than the untransformed controls (Figure 2A-D).

**Figure 2.**
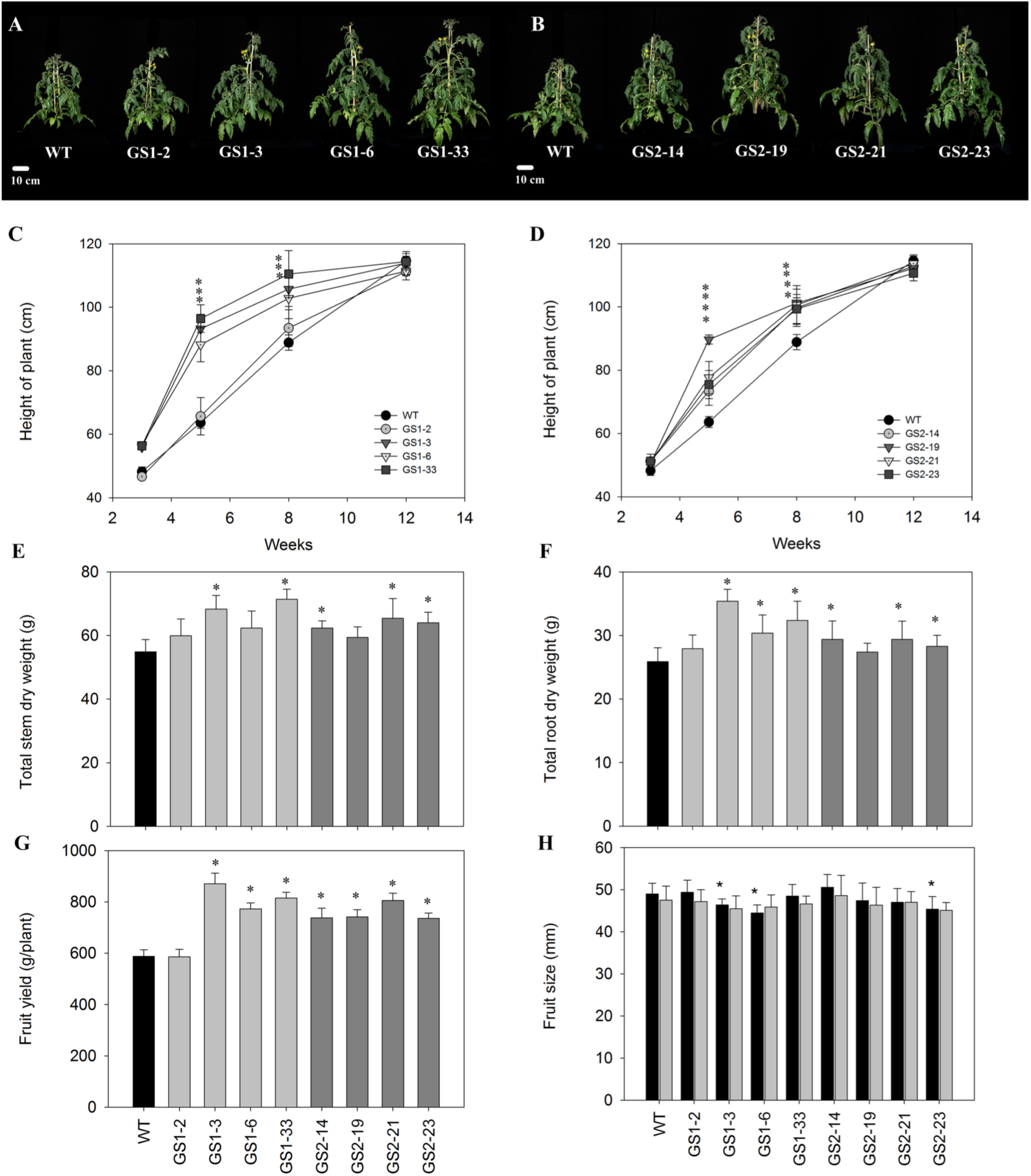
Phenotypic characterization of the *RbcS–GS1* and *RbcS–GS2* lines. **(A, B)** Pictures of 5-week old plants showed faster growth and early flowering. **(C, D)** Plant height along plant development. **(E, F)** Total stem and root dry weight in plants of 16 weeks, respectively. **(G)** Total fruit yield. **(H)** Total fruit size. Values are shown as means ± SE of five individual plants. Asterisks indicate statistically significant differences determined by Student’s *t* test (*P*<0.05).

Both sets of transformants consistently exhibited early flowering (Supplemental Figure 2). As would perhaps be anticipated, this corresponded to an earlier appearance of the first red fruit (Supplemental Figure 2), although the total period of fruit ripening was unaltered.

Despite the observed difference in early growth rate, after 12 weeks the transgenic lines showed no differences from control in plant height. However, at this plant developmental stage, some lines of the transgenic plants displayed an increase in total stem dry weight (GS1-3, GS1-33, GS2-14, GS2-21 and GS2- 23) and increased total root dry weight (GS1-3, GS1-6, GS1-33, GS2-14, GS2- 21, GS2-23) (Figure 2E-F). Interestingly, *RbcS-COX4ts-GS1* and *RbcS- COX4ts-GS2* lines were characterized by a significant increase in fruit yield (Figure 2G), which was consequence of an increase in number of fruits rather than fruit size or weight which were invariant across the lines (Figure 2H). Finally, mature fruits from both sets of transformants showed no significant changes in soluble solid content (°Brix; 3.2 ± 0.2; 3.3 ± 0.1; 3.3 ± 0.1; 3.2 ± 0.3; 3.1 ± 0.2; 3.1 ± 0.2; 3.1 ± 0.1; 3.2 ± 0.2; 3.3 ± 0.2; 3.0 ± 0.2 for control, GS1-2, GS1-3, GS1-6, GS1-33, and GS2-14, GS2-19, GS2-21, GS2-23, respectively), however, a reduction in acidity was observed in the mature fruits of the *RbcS- COX4ts-GS1*. The reproducibility of the results was checked by growing the transgenic plants in two seasons - spring and autumn. With the results from the Autumn season being highly comparable to those presented here (Supplemental Figure 3).

### Mitochondrial over-expression of *GS1* and *GS2* does not impact photosynthesis

Given the observed alteration in plant growth, we next evaluated gas exchange parameters in 5-week-old plants at different photon flux densities (PFD) (Supplemental Figure 4). Interestingly, we did not observe significant changes in net photosynthesis (*A*) or stomatal conductance (*g_s_*) (Supplemental Figure 4A-D); however, we found a slight decrease of maximum PSII photochemical efficiency (*Fv/Fm*) in lines GS1-6, and GS2-23. In view of these changes, we analyzed the electron transport rate (ETR) at different irradiances, since ETR is associated to the flux of electrons passing through PSII in photosynthesis (Tezara et al., 2003). Transgenic lines were characterized by unaltered ETR values measured at low (100 µmol m^-2^) and high (700 µmol m^-2^) irradiances (Supplemental Figure 4E-F). These combined results suggest that overexpression of mitochondrially-targeted *GS1* and *GS2* did not greatly influence the photosynthetic process.

### The effect of over-expression of mitochondrially-targeted *GS1* or *GS2* on leaf starch and pigments biosynthesis and plastidial redox status

To understand further the impact of overexpressing mitochondrially-targeted *GS1* and *GS2* in leaves, we examined starch and pigments levels in fully expanded leaves. Interestingly, we found a reduction of between 25% and 44% in starch levels (with only line GS1-2 exhibiting invariant levels of starch) (Table 1). Additionally, when we measured pigment levels in leaves, we observed significant increase of chlorophyll a (GS1-3, GS1-33, GS2-14, and GS2-19) and chlorophyll b (GS1-3, GS2-14, and GS2-23; Table 2). Finally, by analyzing pyrimidine nucleotide levels, we noted a significant increase in the levels of the reduced pyridine nucleotide NADPH (except in line GS1-2), with an increase in the deduced NADPH/NADP^+^ ratio in the transgenics. However, no changes were noticed in the other pyridine nucleotides, with the exception of the slight decrease in NAD^+^ levels in GS1-3 and GS2-21 lines (Table 1).

**Table 1.** Starch and pyridine nucleotide levels in *RbcS–COX4ts-GS1* and *RbcS–COX4ts-GS2* lines. Leaves were harvested in the middle day. Values are means from five biological replicates ± standard error. Statistically significant differences by Student’s *t* test are indicated by boldface (*P*<0.05). DW, Dry weight.

| Parameters | WT | GS1-2 | GS1-3 | GS1-6 | GS1-33 | GS2-14 | GS2-19 | GS2-21 | GS2-23 |
| --- | --- | --- | --- | --- | --- | --- | --- | --- | --- |
| | $\mu\text{mol/g DW}$ | | | | | | | | |
| Starch | 7.14 ± 0.76 | 6.62 ± 1.04 | **4.17 ± 0.86** | **5.17 ± 0.94** | **4.01 ± 1.14** | **5.06 ± 0.75** | **5.55 ± 0.83** | **5.15 ± 0.93** | **4.75 ± 1.03** |
| NAD<sup>+</sup> | 21.50 ± 0.41 | 20.78 ± 0.35 | 19.86 ± 0.59 | 22.16 ± 0.38 | 21.04 ± 0.30 | 22.06 ± 0.36 | 21.89 ± 0.31 | **20.01 ± 0.29** | 22.78 ± 0.39 |
| NADH | 34.76 ± 1.16 | 33.67 ± 1.07 | **30.26 ± 0.96** | 32.17 ± 1.11 | 31.78 ± 1.28 | 32.67 ± 0.99 | 31.76 ± 1.39 | 33.77 ± 1.36 | 34.03 ± 0.98 |
| NADP<sup>+</sup> | 13.06 ± 0.30 | 12.83 ± 0.27 | 13.87 ± 0.31 | 12.69 ± 0.26 | 13.45 ± 0.33 | 11.97 ± 0.46 | 12.03 ± 0.37 | 13.87 ± 0.33 | 12.98 ± 0.40 |
| NADPH | 5.33 ± 0.25 | 5.82 ± 0.29 | **7.89 ± 0.26** | **7.03 ± 0.18** | **6.78 ± 0.17** | **6.87 ± 0.26** | **6.27 ± 0.19** | **7.02 ± 0.24** | **6.85 ± 0.22** |
| NADH/NAD<sup>+</sup> | 1.86 ± 0.10 | 1.73 ± 0.07 | 1.80 ± 0.06 | 1.58 ± 0.11 | **1.56 ± 0.08** | 1.59 ± 0.15 | **1.53 ± 0.11** | 1.74 ± 0.14 | 1.80 ± 0.10 |
| NADPH/NADP<sup>+</sup> | 0.41 ± 0.04 | 0.47 ± 0.06 | **0.62 ± 0.07** | **0.54 ± 0.06** | **0.53 ± 0.05** | **0.53 ± 0.05** | **0.50 ± 0.03** | **0.56 ± 0.04** | **0.54 ± 0.06** |

**Table 2.** Pigment contents in *RbcS–COX4ts-GS1* and *RbcS–COX4ts-GS2* lines. Data are presented as means ± SE of five biological determinations. Values that are significantly different by Student’s *t* test are indicated by boldface (*P*<0.05). DW, Dry weight.

| Pigments | WT | GS1-2 | GS1-3 | GS1-6 | GS1-33 | GS2-14 | GS2-19 | GS2-21 | GS2-23 |
| --- | --- | --- | --- | --- | --- | --- | --- | --- | --- |
| | $\mu\text{mol g}^{-1}$ DW | | | | | | | | |
| Chlorophyll a | 1.55 ± 0.05 | 1.57 ± 0.07 | **1.75 ± 0.04** | 1.60 ± 0.05 | **1.68 ± 0.03** | **1.66 ± 0.06** | **1.64 ± 0.04** | 1.57 ± 0.06 | 1.59 ± 0.05 |
| Chlorophyll b | 0.54 ± 0.04 | 0.59 ± 0.03 | **0.68 ± 0.03** | 0.58 ± 0.05 | 0.70 ± 0.03 | **0.62 ± 0.03** | 0.59 ± 0.04 | 0.57 ± 0.05 | **0.63 ± 0.03** |
| β-Carotene | 0.034 ± 0.005 | 0.029 ± 0.006 | 0.032 ± 0.005 | 0.030 ± 0.004 | 0.028 ± 0.005 | 0.031 ± 0.004 | 0.033 ± 0.005 | 0.027 ± 0.005 | 0.035 ± 0.004 |

### Metabolic alteration in leaves and their exudates in the transgenic lines and their wild type control

Leaves harvested in the middle of the day were analyzed using an established GC-MS-based metabolite profiling protocol (Fernie et al., 2004) (Figure 3 and Supplemental Data Set 2). The transgenic lines from both constructs displayed similar changes being characterized by decreases in the main hexoses, glucose (lines GS1-3, GS1-6, and GS1-33) and fructose (lines GS1-3. GS1-33, GS2-19, GS2-21) as well as in their phosphorylated derivatives, glucose-6-phosphate and fructose-6-phosphate. Similar changes were displayed by the glucose oxidation products, raffinose (all *RbcS-COX4ts-GS1* lines and GS2-14, GS2-19 lines), and by sucrose (lines GS1-3 and GS2-21). Similarly decreases in trehalose (lines GS1-3, GS1-6, GS1-33, and all *RbcS-COX4ts-GS2* lines), isomaltose (all *RbcS-COX4ts-GS1* lines and GS2-19, GS2-21, GS2-23 lines), 1- *O*-methyl-glucopyranoside (lines GS1-2, GS1-3, GS1-6), and *myo*-inositol (lines GS1-3, GS1-33, GS2-14, GS2-21), were observed. Regarding the organic acids, decreases levels of several tricarboxylic acid cycle intermediates, namely succinate (all lines), fumarate (all *RbcS-COX4ts-GS1* lines and GS2-14, GS2- 21, GS2-23 lines), and malate (all *RbcS-COX4ts-GS1* lines and GS2-14, GS2- 21, GS2-23 lines) were observed, whereas citrate and 2-oxoglutarate levels increased in all transgenic lines. In addition, significant decreases were also observed in caffeoylquinic acid (lines GS1-3 and GS1-33) and quinic acid (lines GS1-2, GS1-3 and GS1-33). Interestingly, some amino acids detected here displayed contrasting behavior including asparagine (all *RbcS-COX4ts-GS1* lines and GS2-14, GS2-21, GS2-23 lines), ß-alanine (all *RbcS-COX4ts-GS1* lines), valine (all *RbcSCOX4ts--GS1* lines), aspartate (all *RbcS-COX4ts-GS1* lines and GS2-14, GS21, GS2-23 lines), arginine (lines GS1-2, GS1-3, GS1-33, GS2-19, GS2-21, GS2-23), *O-*acetyl-serine (GS1-2, GS1-3, GS1-33 lines and all *RbcS-COX4ts-GS2* lines), 4-hydroxyproline (lines GS1-3 and GS2-14), and GABA (all *RbcS-COX4ts-GS1* lines), which all decreased in the transgenic lines. By contrast, serine (all lines), glycine (lines GS1-2, GS1-3, GS1-33, GS2- 14, GS2-19, GS2-23), glutamine (all lines), proline (all lines) and glutamate (all lines) increased. Moreover, the closely related polyamine putrescine was decreased in all *RbcS-COX4ts-GS1* lines as well as lines GS2-19, GS2-21, GS2-23. When taken together these metabolic changes indicate a dramatic shift in primary metabolism with alterations in the levels of sugars, organic acids and also amino acids.

**Figure 3.**
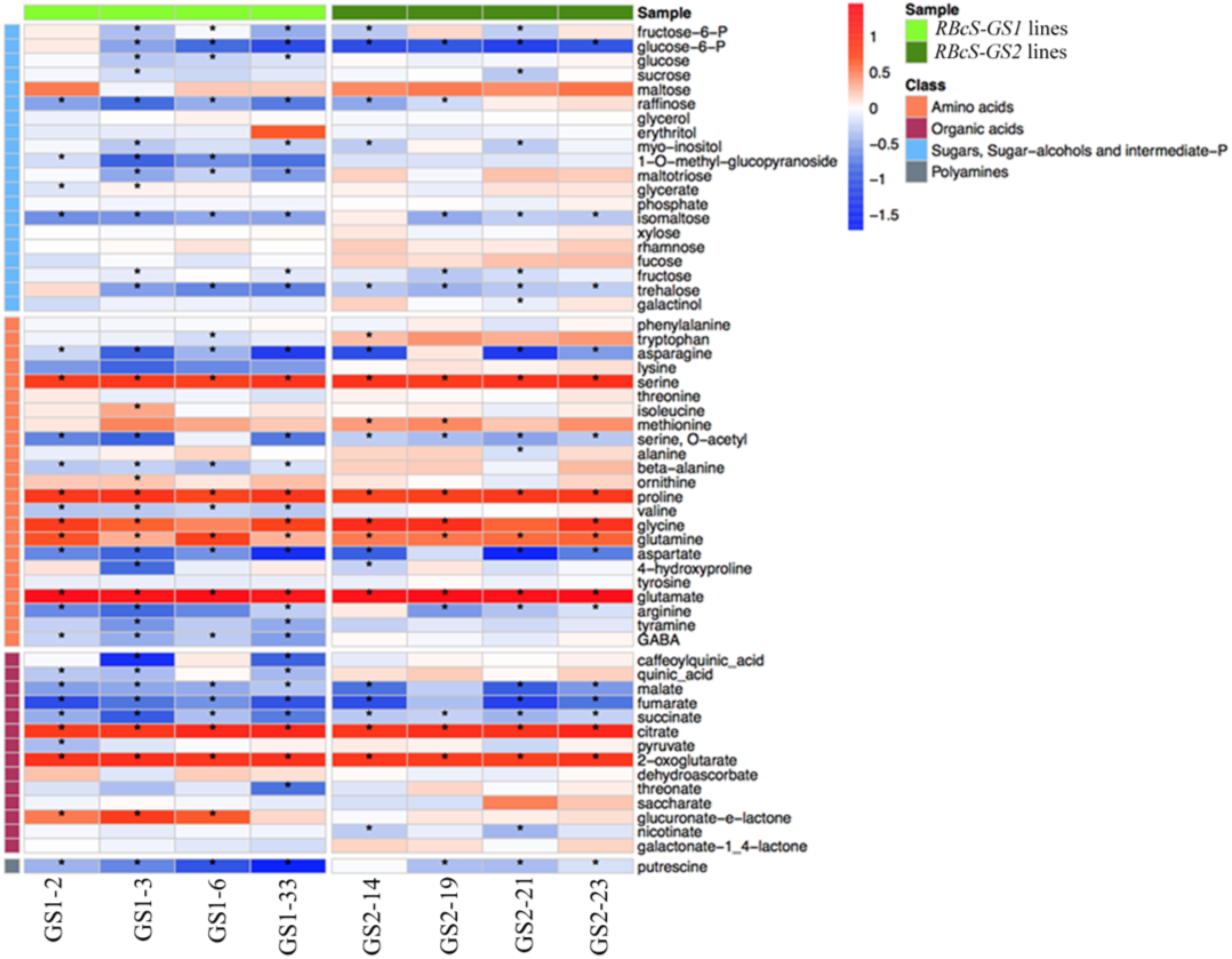
Heatmap of leave primary metabolites in *RbcS–GS1* and *RbcS–GS2* lines. Data were normalized to dry weight and presented as ratio between transgenic plants and WT. Asterisks indicate statistically significant differences determined by Student ’s *t* test (*P*<0.05). The colours indicate the proportional content of each metabolite intensity which has been log10-transformed and mean-centred.

We next evaluated metabolite levels in exudates from leaves of transgenic and control plants. Here, we used authentic standards to perform absolute quantification of metabolite amount (Supplemental Data Set 3). Exudates from all transgenic lines were characterized for a large increase in the major sugars, glucose, fructose, and sucrose (up to 6.5-fold). Changes were also apparent in glutamate which decreased in the exudates from all transgenic lines and GABA levels in the exudates from line GS1-3, GS1-33, GS2-14 and GS2-21.

### Effects of over-expression of mitochondrially-targeted *GS1* or *GS2* on enzyme activities concerned with nitrogen assimilation and other aspects of primary metabolism

The aforementioned results indicate that plants over-expressing *GS1* and *GS2* targeted to the mitochondria exhibit reduced starch biosynthesis in source leaves. The plastidic phosphoglucomutase (PGM) and ADP-glucose pyrophosphorylase (AGPase) activities have been described as key enzymes for starch biosynthesis and its regulation (Caspar et al., 1985; Hanson and Mchale, 1988; Lin et al., 1988; MacRae and Lunn, 2006). In addition, it has been previously demonstrated in potato that phospho*enol*pyruvate carboxylase (PEPC) activity also plays an important role in starch metabolism (Rademacher et al., 2002). For this reason, we analyzed the PGM, PEPC and AGPase activities in leaves of the transgenic and control lines. This analysis revealed that both the *RbcS-COX4ts-GS1* and the *RbcS-COX4ts-GS2* lines displayed significant decreases in PGM and AGPase enzyme activities, whereas PEPC activity was invariant in the transformants (Table 3). In addition, we observed an increase in total and initial activities of NADP-malate dehydrogenase (NADP- MDH) of the chloroplast. These data, when taken together, suggest an altered C metabolism in chloroplast. By contrast, in considering the tricarboxylic acid (TCA) cycle, there was a lack of consistent changes in NAD-MDH and fumarase activities (Table 3).

**Table 3.** Enzymatic activities in *RbcS–COX4ts-GS1* and *RbcS–COX4ts-GS2* lines. Leaves were harvested at middle day. Values are means from five biological replicates ± standard error. Statistically significant differences by Student’s *t* test are indicated by boldface (*P*<0.05). Phosphoglucomutase (PGM), ADP-glucose pyrophosphorylase (AGPase), nitrate reductase (NR), nitrite reductase (NiR), glutamate dehydrogenase (GDH), glutamate synthase (GOGAT). DW, Dry weight.

| Enzymes | WT | GS1-2 | GS1-3 | GS1-6 | GS1-33 | GS2-14 | GS2-19 | GS2-21 | GS2-23 |
| --- | --- | --- | --- | --- | --- | --- | --- | --- | --- |
|  | nmol min<sup>-1</sup> g<sup>-1</sup> DW |  |  |  |  |  |  |  |  |
| PGM | 1868 ± 126 | 1768 ± 156 | **1257 ± 117** | **1397 ± 137** | **1308 ± 120** | 1598 ± 129 | **1497 ± 110** | **1427 ± 147** | **1528 ± 132** |
| AGPase | 1468 ± 86 | 1387 ± 69 | **1186 ± 65** | **1287 ± 76** | **1157 ± 59** | **1298 ± 79** | **1305 ± 69** | **1228 ± 72** | **1176 ± 63** |
| PEPC | 176 ± 12.6 | 187 ± 9.3 | 160 ± 8.8 | 164 ± 10.3 | 172 ± 12.1 | 164 ± 8.4 | 183 ± 10.4 | 170 ± 7.6 | 193 ± 11.5 |
| NADP-MDH total | 77.4 ± 5.2 | 83.4 ± 7.2 | **104.3 ± 6.4** | **96.3 ± 6.3** | **117.3 ± 7.5** | **88.2 ± 5.3** | **92.4 ± 6.4** | **93.2 ± 6.3** | **102.3 ± 4.7** |
| NADP-MDH initial | 28.3 ± 2.7 | 26.3 ± 1.8 | **37.7 ± 2.1** | **34.2 ± 1.8** | **41.4 ± 3.2** | **33.7 ± 1.7** | **36.9 ± 2.4** | **39.3 ± 1.3** | **34.6 ± 1.8** |
| NADP-MDH activation stage | 11.7 ± 1.4 | 10.4 ± 1.7 | 11.3 ± 1.4 | 10.2 ± 2.1 | 9.8 ± 2.1 | 10.3 ± 1.8 | 11.3 ± 1.6 | 12.3 ± 2.4 | 12.1 ± 1.7 |
| NAD-MDH (mmol min<sup>-1</sup> g<sup>-1</sup> DW) | 18.3 ± 2.7 | 16.5 ± 3.9 | 17.2 ± 2.6 | 18.5 ± 3.2 | **22.3 ± 3.2** | 19.4 ± 2.6 | 18.9 ± 3.5 | **20.4 ± 1.7** | 16.4 ± 3.5 |
| Phosphoenolpyruvate carboxylase | 88.3 ± 4.8 | 84.2 ± 4.7 | 89.4 ± 5.3 | 94.3 ± 4.5 | 92.2 ± 4.3 | 83.4 ± 5.9 | 95.2 ± 4.2 | 86.3 ± 6.2 | **80.3 ± 3.1** |
| Fumarase | 1075 ± 82 | 1159 ± 92 | 1046 ± 103 | 1143 ± 79 | 1064 ± 72 | 1173 ± 93 | 1032 ± 103 | 1064 ± 94 | 1052 ± 83 |
| NR | 11.6 ± 4.6 | 15.8 ± 3.5 | **21.4 ± 4.1** | **18.3 ± 3.2** | **17.4 ± 3.1** | **18.3 ± 3.3** | **19.3 ± 4.2** | **20.2 ± 3.1** | **17.3 ± 4.6** |
| NiR (U min<sup>-1</sup> g<sup>-1</sup> DW) | 2.1 ± 0.6 | 1.9 ± 0.8 | **1.3 ± 0.4** | **1.1 ± 0.5** | **0.9 ± 0.2** | **1.6 ± 0.4** | **1.3 ± 0.2** | **1.5 ± 0.4** | **1.3 ± 0.2** |
| GDH | 26.4 ± 2.1 | **32.4 ± 1.5** | **42.3 ± 2.5** | **39.3 ± 2.1** | **44.5 ± 3.2** | **32.5 ± 1.8** | **29.4 ± 2.6** | **31.4 ± 3.1** | **28.6 ± 1.6** |
| GOGAT | 5.3 ± 1.3 | 4.7 ± 1.4 | **3.2 ± 0.8** | **3.8 ± 0.8** | **2.4 ± 0.9** | 4.2 ± 0.7 | **3.3 ± 0.8** | **3.6 ± 1.1** | **4.0 ± 0.8** |

Given the amino acids profile obtained and shown in Figure 3, we examined if leaf-specific mitochondrial GS expression affected N metabolism at the level of enzyme activities. We evaluated the activities of the main N assimilation-related enzymes. Interestingly, the nitrate reductase (NR) activity was increased in transgenic lines whereas the nitrite reductase (NiR) decreased. Moreover, glutamate dehydrogenase (GDH) activity was significantly higher in all transgenic lines when compared to the control; however, the glutamate synthase (GOGAT) activity of the transgenic lines was reduced in comparison to that observed in the control leaves (Table 3). These results suggest considerable shift in N metabolism in the transgenic leaves. Taking this observation into account, alongside the altered glycine/serine levels displayed by the transgenic lines (Figure 3), we explored the photorespiratory interconnection between N and C metabolism in these lines. In mitochondria, one of the key reactions of photorespiration is catalyzed by the glycine decarboxylase complex (GDC), which catalyzes the reaction converting two molecules of glycine in one molecule of serine generating CO_2_ and NH_4_^+^ (Bauwe et al., 2010). GDC is a hetero-tetramer formed by four enzymes (P-, T-, H-, and L-) and their combined activity has been described as the major control point for photorespiratory flux being subject to multi-level regulation (Timm, 2020). For instance, mitochondrial redox imbalance affects GDC activity through the GDC L-protein (Timm, 2020). Interestingly, in this respect, the expression level of the GDC L-protein was significantly higher in all transgenic lines in comparison to the control line (Supplemental Figure 5).

### Metabolic alterations during development and ripening of the RbcS- COX4ts-GS1 and RbcS-COX4ts-GS2 fruits

The changes in primary metabolism during fruit development and ripening in the transformants were analyzed by using the same protocol as for leaves. We evaluated the metabolite levels in fruits harvested 22 d after pollination (DAP), and at mature green (MG), and ripe (R) stages. In general, similar patterns of changes were observed when comparing the RbcS-COX4ts-GS1 and RbcS- COX4ts-GS2 lines. At the ripe stage, both lines exhibited an increase in most of the free amino acids, including serine, threonine, alanine, ß-alanine, proline, valine, glycine, aspartate in comparison to control fruits (Figure 4, 5). However, the levels of glutamine, glutamate, tyrosine, GABA, and methionine significantly were significantly lower in the transgenic lines at the ripe stage. We additionally observed significant increases in the levels of sucrose at 22 DAP and in ripe fruit (between 1.2 and 1.4-fold of wild type levels at 22 DAP and between 1.8 and 2.2-fold of wild type levels at the ripe stage), although no significant changes were observed in the GS1-2 line. Similarly, an increase in the levels of the sugar phosphates, glucose-6P and fructose-6P, was observed in all lines. Interestingly, these changes were coupled with an increase in the level of starch at 22 DAP stage in all transgenic lines (235.2 ± 18.5; 255.2 ± 15.3; 387 ± 20.4; 315.2 ± 11.8; 375.4 ± 15.8; 336.7 ± 21.4; 367.2 ± 17.3; 299.4 ± 12.1; 287.4 ± 16.4 nmol g^-1^ DW for control, GS1-2, GS1-3, GS1-6, GS1-33, and GS2-14, GS2-19, GS2-21, GS2-23, respectively). Two disaccharides associated to starch degradation, maltose and isomaltose, displayed slightly increased levels at the 22 DAP stage in all lines while isomaltose levels were lower at mature green and ripe stages in the transgenics in comparison to the control (Figure 4, 5). These changes were accompanied by increases in the other major sugars namely fructose and glucose, that also increased at ripe stage. By contrast, the levels of saccharate and raffinose, products released following glucose oxidation, were lower at the ripe stage in the transgenic lines (Figure 4, 5). Perhaps surprisingly given that significant differences were found in the levels of major sugars in ripe fruits, we did not observe significant differences in the total soluble solids content. Despite the levels of fumarate and malate being invariant in the fruit of the transgenics other metabolic changes of note were that citrate was observed at significantly higher levels at the ripe stage in all lines. Furthermore, the levels of the cell wall-related sugars, xylose and rhamnose, which increase during fruit ripening consistent with the known increased cell wall degradation at this time point, were higher in the transgenic lines in comparison to control fruits (Figure 4, 5). Moreover, significantly lower putrescine levels were observed at both 22 DAP and mature green stages but significantly higher levels were observed in ripe fruit in all transgenic lines (Figure 4, 5).

**Figure 4.**
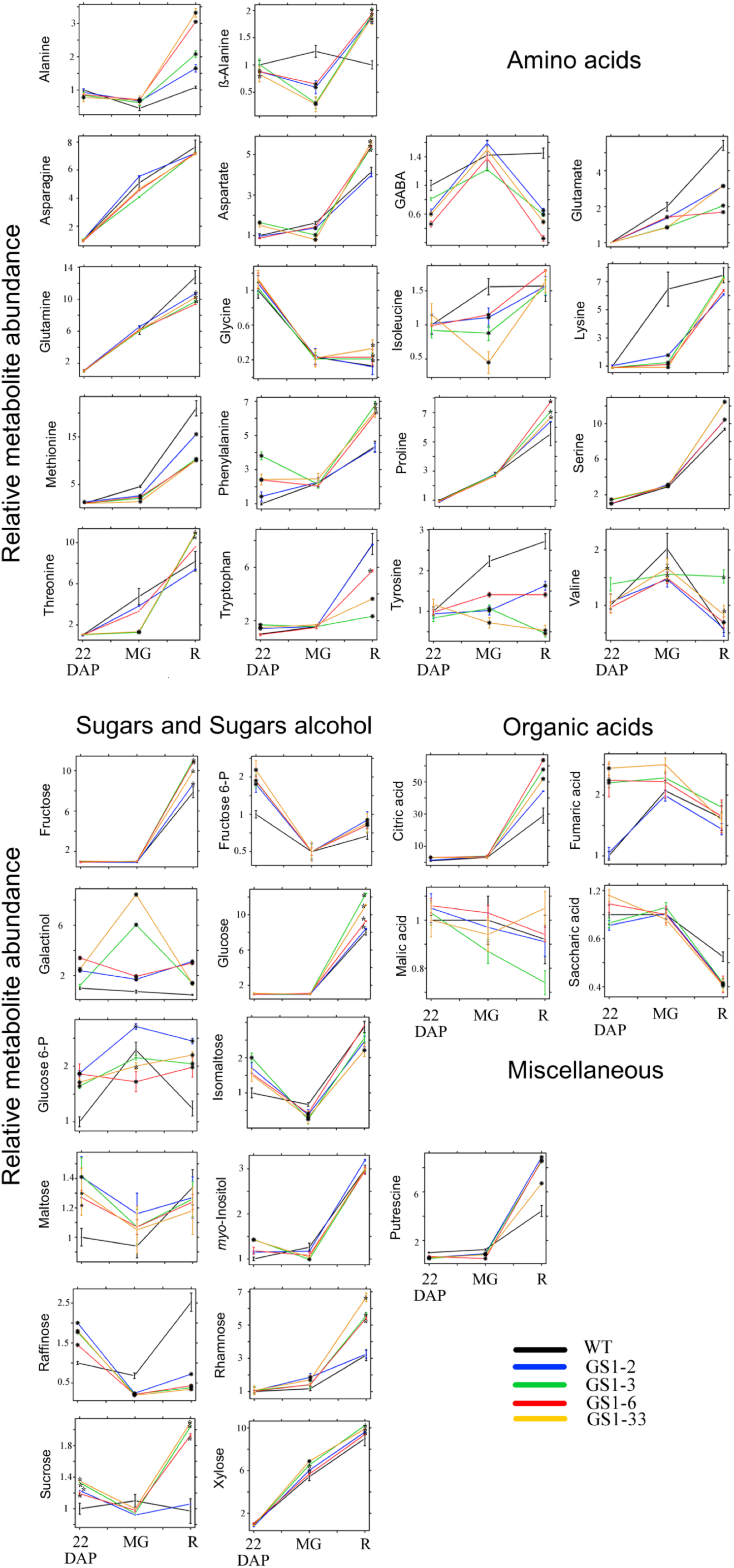
Primary metabolites in WT and *RbcS–GS1* lines at three developmental and ripening stages (22 d after pollination (DAP), mature green (MG), and ripe (R)). Values are normalized to mean value of WT at 22 DAP stage. Data showed as means ± SE of five individual plants. Asterisks indicate statistically significant differences determined by Student ’s *t* test (*P*<0.05) of the transgenic line compared to WT at the same developmental stage.

**Figure 5.**
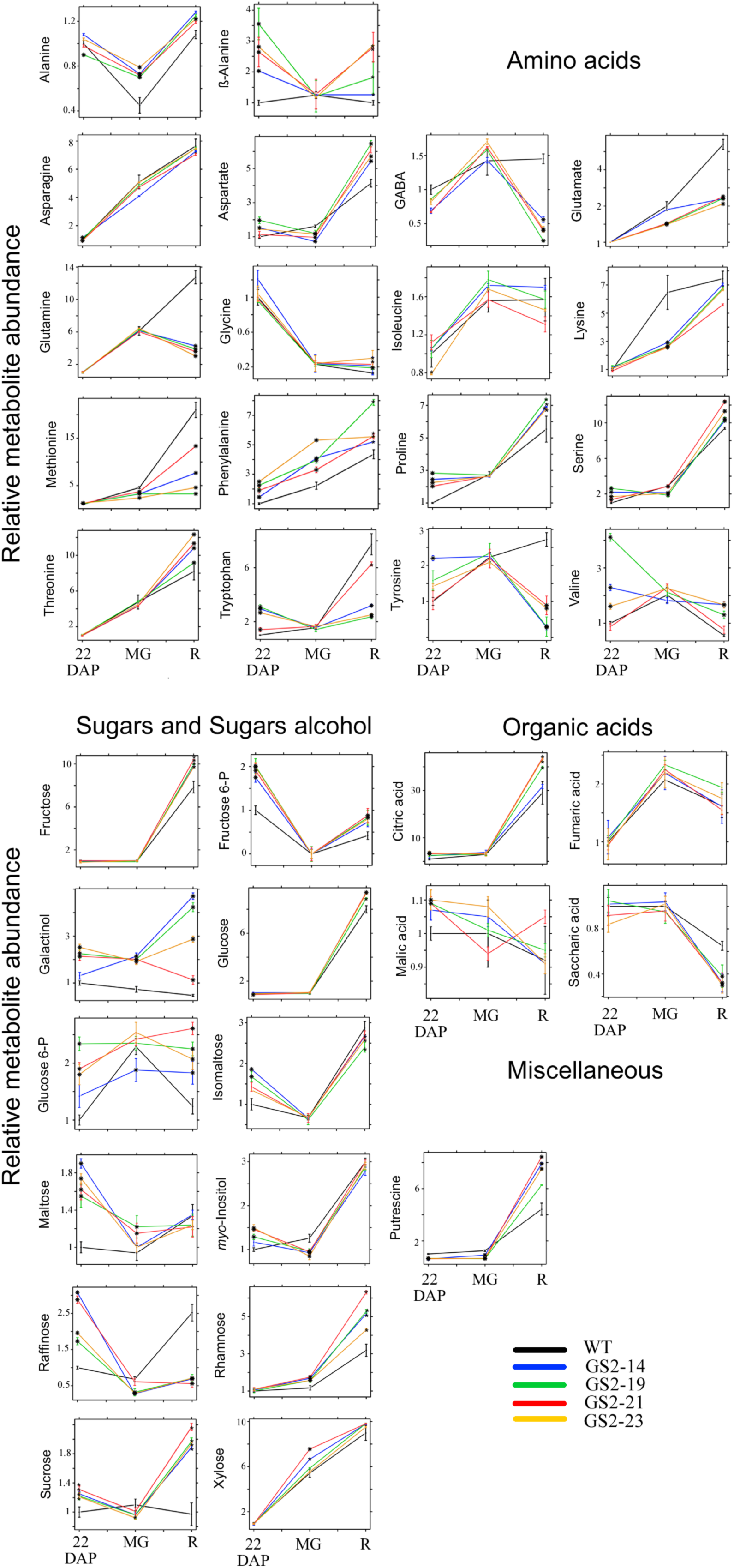
Primary metabolites in WT and *RbcS–GS2* lines at three developmental and ripening stages (22 d after pollination (DAP), mature green (MG), and ripe (R)). Values are normalized to mean value of WT at 22 DAP stage. Data showed as means ± SE of five individual plants. Asterisks indicate statistically significant differences determined by Student ’s *t* test (*P*<0.05) of the transgenic line compared to WT at the same developmental stage.

### Effect of increasing mitochondrial GS activity in leaves on metabolic fluxes in the fruits

Given the changes in sugars and starch levels in small fruits (22 DAP), we next decided to assess the relative rate of carbon fluxes by incubation of fruit pericarp discs at 22 DAP stage of control and two GS1 lines (GS1-3 and GS1- 33) and two GS2 lines (GS2-14 and GS2-19) with positionally-labelled ^14^C- glucose. Samples were incubated in [1-^14^C]-glucose and [3,4-^14^C]-glucose for 5 hours and the ^14^CO_2_ evolution was monitored. The CO_2_ releases from C1 of glucose is associated to respiration through the oxidative pentose phosphate pathways while the respiration through TCA cycle is linked to release from C3,4 position of glucose. Therefore, a high ratio of CO_2_ evolution from C3,4 to C1 position of glucose provides strong evidence of mitochondria activity. When comparing the ^14^CO_2_ releases from control and RbcS-COX4ts-GS1 and RbcS- COX4ts-GS2 lines, we observed a significant increase in the C3:4/C1 ratio in transgenic fruits (Figure 6).

**Figure 6.**
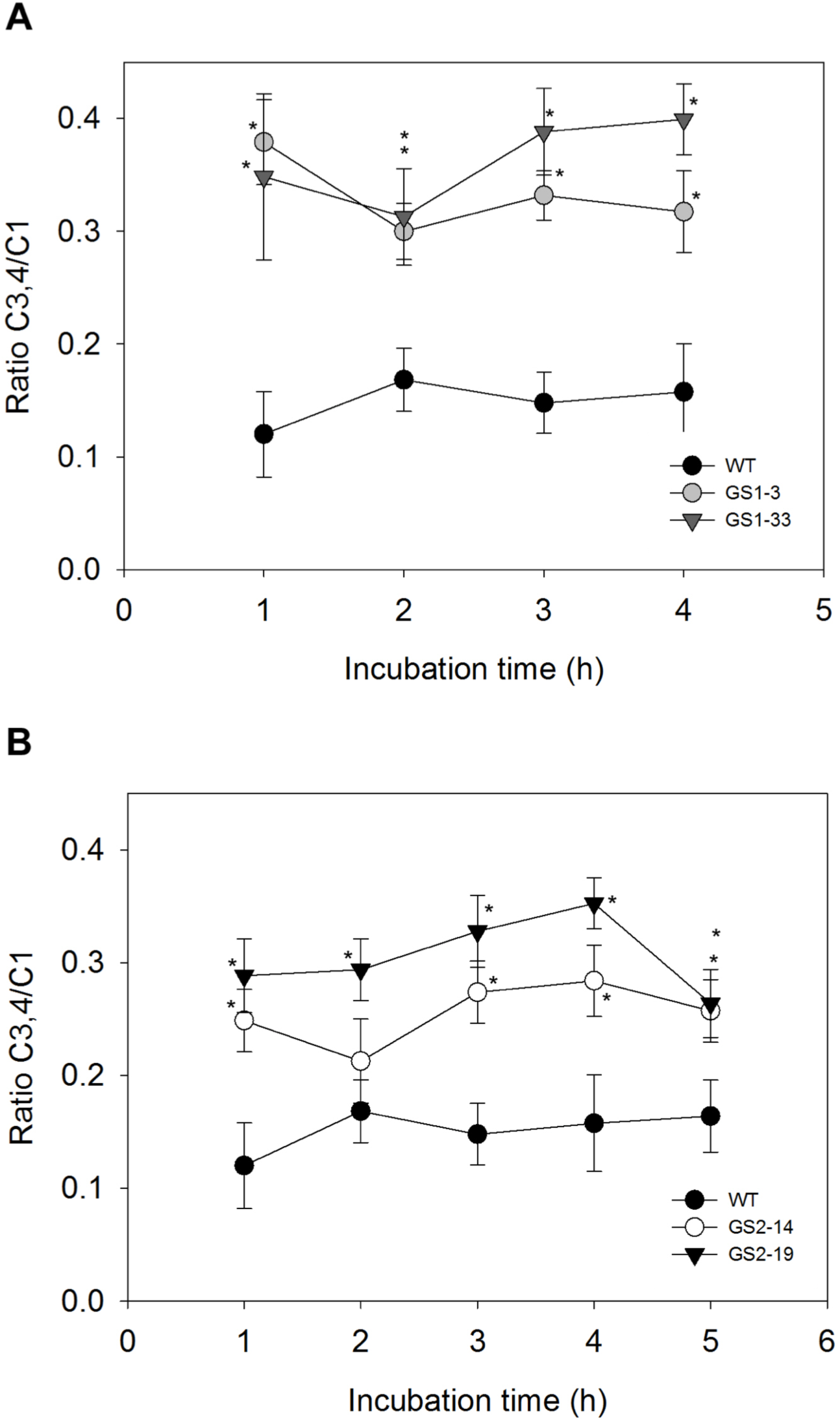
Ratio of CO_2_ evolution from C3.4/C1 positions of glucose in pericarp fruits of 22 DAP stage from *RbcS–GS1* and *RbcS–GS2* lines.

In order to expand this study, same samples were incubated with [U-^14^C]- glucose for 3 hours to assess the redistribution of radiolabel. We did not observed changes in total label incorporated; however, differences were apparent when comparing radiolabel redistribution (Table 4). In agreement with the steady-state metabolite data, significantly increased label incorporation into starch and the total hexose phosphate pool was observed in the analyzed lines at the 22 DAP stage (lines GS1-3, GS1-33, GS2-14, and GS2-19). However, no significant changes were observed in either label redistribution to cell wall, organic acids, amino acids or sucrose (Table 4). This pattern was also seen following estimation of the fluxes with an elevated rate of starch synthesis being the only consistent change, although it is important to note that the rate of amino acid biosynthesis was also observed to slightly decrease in line GS1-3.

**Table 4.** Changes in respiratory carbon redistribution of radiolabel carbon after incubation of ^14^C- Glc in pericarp discs from RbcS-COX4ts-GS1 and RbcS-COX4ts-GS2 lines at 22 DAP stage.

| Parameter | WT | GS1-3 | GS1-33 | GS2-14 | GS2-19 |
| --- | --- | --- | --- | --- | --- |
| Label incorporated (Bq) |  |  |  |  |  |
| Total uptake | 380.8 ± 34.5 | 382.3 ± 23.8 | 397.5 ± 47.3 | 379.3 ± 38.2 | 401.4 ± 38.6 |
| Cell wall | 8.5 ± 0.6 | 8.1 ± 0.9 | 7.7 ± 0.7 | 9.1 ± 1.0 | 8.6 ± 0.8 |
| Starch | 55.6 ± 7.7 | **72.6 ± 9.5** | **78.2 ± 7.5** | **66.6 ± 4.6** | **70.5 ± 6.6** |
| Organic acids | 150.6 ± 11.7 | 132.5 ± 8.7 | 137.4 ± 9.3 | 137.3 ± 10.3 | 150.5 ± 9.3 |
| Total hexose phosphate | 40.7 ± 9.4 | **55.8 ± 5.3** | **52.4 ± 6.3** | **64.5 ± 8.3** | 46.3 ± 9.3 |
| Amino acids | 88.6 ± 7.3 | 79.6 ± 9.5 | 93.5 ± 8.7 | 85.7 ± 6.9 | 90.6 ± 11.5 |
| Suc | 33.6 ± 4.8 | 30.6 ± 6.7 | 25.5 ± 5.7 | 23.7 ± 7.9 | 32.7 ± 6.9 |
| Metabolic flux (nmol g<sup>-1</sup> DW h<sup>-1</sup>) |  |  |  |  |  |
| Cell wall | 85.6 ± 7.5 | 77.6 ± 8.3 | 91.7 ± 9.2 | 80.7 ± 7.7 | 95.8 ± 8.4 |
| Starch | 167.8 ± 11.7 | **247.4 ± 20.6** | **210.4 ± 15.3** | **196.7 ± 9.4** | **216.5 ± 11.5** |
| Organic acids | 306.7 ± 18.7 | 275.5 ± 23.7 | 269.4 ± 22.7 | 315.6 ± 16.3 | 289.4 ± 24.3 |
| Amino acids | 278.3 ± 12.4 | **238.4 ± 9.5** | 262.6 ± 13.5 | 296.6 ± 13.5 | 275.5 ± 7.8 |
| Suc | 57.3 ± 8.4 | 68.3 ± 12.5 | 50.7 ± 9.7 | 73.7 ± 7.7 | 63.4 ± 8.3 |

## DISCUSSION

During recent decades, considerable effort has been dedicated to elucidate molecular mechanisms associated with enhanced biomass. N and C metabolism are tightly coordinated in the fundamental biochemical pathways that support plant growth (Nunes-Nesi et al., 2010; Perchlik and Tegeder, 2018; Lu et al., 2020). Understanding of the regulation of these pathways is therefore essential for optimizing plant growth in a manner that can improving crop production (Sonnewald and Fernie, 2018; Fernie et al., 2020; Sonnewald et al., 2020). As such many different studies have focused on N metabolism and C/N interactions (Perchlik and Tegeder, 2018; Lu et al., 2020; Vallarino et al., 2020), with a couple of these attempting to use multi-gene approaches in order to influence both C and N metabolism simultaneously (Zhang et al., 2015; Vallarino et al., 2020). Here, we postulate, as previously described, that plant metabolic pathways change to be adaptable to combat different environment conditions (Bohnert et al., 1995), although sometimes this adaptation results in yield losses (Amthor et al., 2019). In this study, we used constraint scanning of an FBA leaf model to predict metabolic flux distributions associated with an increase in metabolic efficiency in leaves. The simulations showed a potential mechanism to improve leaf metabolic efficiency and increase biomass with reduced energy costs was to move a functional glutamine synthetase (GS) into the mitochondria. GS catalyzes the ATP dependent biosynthesis of Gln from Glu and NH_4_^+^. There are two GS isoforms, a cytoplasmic form (GS1) and a plastidic form (GS2). GS2 is described to be involved in the re-assimilation of NH_4_^+^ produced during photorespiration and nitrate reduction while GS1 has been associated to the assimilation of NH_4_^+^ produced from other physiological processes (Seger et al., 2015). Moreover, a study in Arabidopsis showed that GS2 isoform can be dual targeted to leaf plastid and mitochondrial (Taira et al., 2004). The authors reported induced leaf mitochondrial GS activity in response to either CO_2_ limitation or transient darkness (Taira et al., 2004). Also, the SUBA4 database suggested that there is also proteomic evidence for the presence of GS2 in Arabidopsis mitochondria (Hooper et al., 2017).

Here, we evaluated the effect of independently overexpressing the two GS isoforms in mitochondria under the control of the leaf mesophyll-specific promoter of ribulose-bisphosphate carboxylase (RbcS). This allowed us to assess whether mitochondrial manipulation of GS activity produced the physiological changes in tomato plants that we anticipate from the model. The results obtained indicated that modified mitochondrial GS activity can indeed result in altered plant development due to elevated source to sink transport. The observed changes included increased and earlier flowering and fruit set which culminated in both a shortened life cycle and an enhanced yield.

### Effects of overexpression of a mitochondrially-targeted GS on leaf metabolism

A considerable amount of research has long suggested that N assimilation and remobilization are important interconnected processes which play key roles in plant growth and development (Miflin and Habash, 2002). Here, we provide experimental evidence that the mitochondrially-targeted overexpression of *GS1* or *GS2* in mitochondria, led to similar alterations in plant growth and development. When taken together, our data suggest that shifting GS activity to the mitochondria, renders leaves more efficient with regard to their N assimilation/mobilization pathways.

Irrespective of the gene used the increased GS activity in mitochondria of the transgenics resulted in an enhanced rate of early plant growth with plants also producing more fruits. This phenomenon demonstrates that shifting the GS activity to the mitochondria can improve the efficiency of leaf metabolism, thereby improving the source strength of the leaves. This observation was in concordance with higher values of shoot and root biomass together with a notable rise in the content of major amino acids and total soluble proteins in leaves (which represent the major source of organic nitrogen in plants).

Interestingly, despite displaying apparently elevated source strength, the transgenic lines did not display major changes in photosynthetic parameters - yet were characterized by lower levels of soluble sugars and starch than the wild type control. This finding indicates that the transgenics are likely characterized by a higher C export to sink organs. This raises the likelihood of an increased flux of trioses-Ps from the chloroplast to the cytosol which via previously defined mechanisms (Heldt and Flugge, 1987; Stitt, 1990a), would increase the cytosolic hexoses-P pool and thereby drive a higher rate of Suc synthesis. Consistent with this hypothesis is the fact that we observed higher Suc levels in leaf exudates as well as reduced enzymatic activities of phosphoglucomutase (PGM) and ADP-glucose pyrophosphorylase (AGPase) in the leaves of both sets of transgenic plants. Given that these enzymes have been documented to play key roles in starch biosynthesis (Sweetlove et al., 1999; Fernie et al., 2001), which is essentially competitive to sucrose biosynthesis in the illuminated leaf (Stitt, 1990b), these data fully fit our proposed FBA model results. Furthermore, AGPase activity appears also to be modulated at the post-translational level (Hendriks et al., 2003). One of the regulatory mechanisms coordinating the activity of this enzyme is associated to the plastidial redox state, with malate playing a key role in mediating intra- organellar redox metabolism (Centeno et al., 2011; Szecowka et al., 2012) Interestingly, overexpression of GS1 and GS2 in mitochondrial produced a significant increase in the NADPH level altering the NADPH/NADP^+^ level in photosynthetically active leaves, which is also consistent with altered AGPase activity in these lines. These transgenic plants showed also altered malate levels in leaves. Therefore, the results are strongly in agreement with previous evidence that starch metabolism is modified by redox active intermediates levels including malate (Centeno et al., 2011; Osorio et al., 2013).

In addition to the above-described effect of mitochondrial GS1 or GS2- overexpression, two important observations can be highlighted in the transgenic leaves in comparison to WT: i) an activation in the initial and total activities of the NADP-dependent malate dehydrogenase (NADP-MDH) of the chloroplast – which uses excess NADPH to convert oxaloacetate to malate; and ii) slight increase in photosynthetic pigment accumulation. Both observations have been commonly used as diagnostic marker for alterations in plastidial redox status (Scheibe, 1991; Scheibe et al., 2005; Nashilevitz et al., 2010). However, we cannot exclude that these changes in chlorophyll content can be due to the observed alteration in the level of the amino acid Glu in the leaves of the transformants, since Glu is the precursor for tetrapyrrole ring biosynthesis (Von Wettstein et al., 1995; Lytovchenko et al., 2011).

Our results suggest that the metabolic signaling-network mechanisms that allow C and N to be remobilized from leaves to increase biomass and fruit production by modifying mitochondrial GS activity, may be associated to changes in the redox state in leaves. In this vein, Chaouch et al., (2010) previously suggested that photorespiration plays a key role in the regulation of cellular redox status. Photorespiration, furthermore, interacts with a range of diverse metabolic processes including but not limited to photosynthesis, N metabolism, respiration (Foyer et al., 2009; Bauwe et al., 2010; Bloom et al., 2010; Fernie et al., 2013) and has previously been positively associated to productivity (Aliyev, 2012). In this study, we observed indicators of an increased photorespiratory flux in that we detected a significant increase mRNA levels of the L subunit of the glycine decarboxylase complex (GDC) and an accumulation in Ser levels. The higher rate of photorespiration in the transgenic plants together with the increased NADP-MDH activity described above, is suggestive of activation of the malate valve in order to transport the excess of reducing equivalents out of chloroplast. In keeping with this conclusion are the previous results demonstrating that tobacco leaves displaying reduced expression of NADP-MDH were also characterized as displaying decreased rates of photorespiration (Backhausen and Scheibe, 1999).

Previous studies have also described a close relation between photorespiration and 2-oxoglutarate (2-OG) metabolism (including both the TCA cycle and GABA shunt) (Eisenhut et al., 2013; Carroll et al., 2015; Fromm et al., 2016). Our results here suggest that functions of chloroplasts and mitochondrial compartments are tightly coordinated through redox and intracellular metabolite pool exchanges as previously described (Nunes-Nesi et al., 2005; Noguchi and Yoshida, 2008; Araújo et al., 2012; Osorio et al., 2013). Our results also demonstrated a marked shift in 2-OG metabolism. Evaluation of the steady- stage metabolite levels were, by and large, in keeping with a reduction in flux through the mitochondrial steps of the TCA cycle in that the TCA intermediates succinate, fumarate, and malate were reduced in the leaves with increased mitochondrial GS activity. By contrast, the levels of 2-OG and citrate were dramatically increased. Interestingly, we also noticed altered levels of several amino acids derived from 2-OG such as Glu, Pro, Gln. Although our results suggested a substantial suppression of TCA cycle activity this was not reflected in the maximal catalytic activities of its constituent enzymes. However, this is not overly surprising since previous studies have suggested that these enzymes are present in levels vastly in excess of what is needed in terms of the fluxes they bear (Gibon et al., 2004).

The observed changes in different accumulation in metabolites directly related to N assimilation pathway, especially Glu, Pro, Gln, Asn, Arg, and 2-OG levels prompt the question, how can the TCA cycle provide enough carbon skeletons to increase N assimilation rather than synthesis of ATP?. With the observation of higher citrate levels in all transgenic lines (Figure 3), it could be postulated that this citrate, which includes citrate accumulated in leaf vacuoles during the night, could contribute to the increased C skeleton demand for N assimilation. At the same time, the availability of NADH from glycine decarboxylase flux (photorespiration) should be sufficient to support the mitochondrial electron transport flux required for ATP synthesis. In this study, the activity of N- assimilatory enzymes in transgenic leaves displayed interesting results. As expected we detected higher GS activity, but also, we observed a significant increase in nitrate reductase (NR) and glutamate dehydrogenase (GluDH) while nitrite reductase (NiR) and glutamate synthetase (GOGAT) were decreased in the transgenic plants. Our results thus underline the previously described (Hodges, 2002), strong links between 2-OG metabolism and N assimilation.

### Effects of mitochondrially targeted overexpression of GS on flowering, fruit set, and fruit metabolism and yield

As a sink tissue, fruit strength is largely dependent on C supply from the phloem, especially in the form of Suc (Zrenner et al., 1995; Sonnewald et al., 1997; Fridman et al., 2004; Obiadalla-Ali et al., 2004). Our data indicate that mitochondrial GS activity improves N assimilation and increase photoassimilate partitioning and transport to fruit. This had several agronomically important consequences namely the transgenic tomato plants flowered earlier and more frequently and fruit set was correspondingly also earlier and more frequent. Given that at least at Northern latitudes most tomatoes are grown year-round in greenhouses this in itself could constitute a major gain since it may allow and increased number of harvests per year. The results presented here suggests that despite displaying earliness in growth rate and flowering, the fruit size was similar or even smaller that the wild type. That said, the mitochondrially targeted over-expression of GS result in enhanced overall tomato yield, which suggests that the model predicted a highly useful intervention.

We also observed a higher rate of starch accumulation rate during the fruit growth stage in the transgenic plants that, as we have seen previously (Centeno et al., 2011), corresponded to increased Glc, Fru and Suc levels in ripe fruits. Consistent with these changes in the starch accumulation at the 22 DAP stage from both lines was the increase in the starch biosynthetic flux and, also, increased label into pool of hexose phosphate. These data also support previous observation that starch accumulation during fruit development is an important source of sugars in ripe fruits (Baxter et al., 2005; Osorio et al., 2013; Colombié et al., 2015). It is important to remember here that the genetic intervention was leaf mesophyll specific so the fact that altered flux was maintained in isolated pericarp discs from the transformants suggests that the latency of these differences is caused by a reprogramming of sink metabolism as a consequence to changes in the source. This observation leads us to propose that an elevated sink strength, at least partially drove the higher sugar influx from source leaves into transgenic fruits, increasing the rate of starch synthesis and consequently the soluble sugar levels during fruit ripening. It will be interesting in future studies to address the precise mechanism by which this is achieved.

We additionally observed higher levels of the cell wall associated sugars, xylose and rhamnose, during fruit ripening in both transformants which may indicate higher degradation rate of the cell wall pectin; however, the results from the feeding experiments did not find changes in the total label incorporated to the cell wall. Other metabolic changes found in ripe fruit of the transformants, such as increase of amino acids potentially reflect a higher protein degradation rate at late ripening stages in order to support the increased respiration rate through the TCA found in ripe fruit in both sets of transformants. In addition, a significant accumulation of putrescine was observed in the ripe fruits from both lines. Putrescine is one of the most prevalent polyamines in plants and has been implicated in alkaloid biosynthesis as well as being a mediator of several signaling pathways (Bienz et al., 2005; Mattoo and Handa, 2008). Catabolism of polyamines has been described to feed TCA cycle via the GABA pool (Rastogi and Davies, 1991; Bienz et al., 2005). However, further research is needed in order to ascertain the significance of these changes.

In conclusion, we demonstrate here that overexpressing a targeted GS activity in leaves is an effective strategy for improving leaf metabolic efficiency and both the number of fruit set, the earliness of this developmental transition and ultimately an enhanced fruit yield. In light of growing interest in urban farming (Kwon et al., 2020; Fernie and Yan, 2020) alongside the fact that a large proportion of global tomato growth occurs in greenhouses, the earliness phenotype alone is likely of high agronomic value as the shortening of the life cycle potentially allows a greater number of harvests per year. Our results provide evidence that by moving the GS activity into mitochondria in leaves, we observed a strong effect on the source strength of the leaves which were more efficient in N assimilation/mobilization. It appears that the higher fruit production observed was due to altered photoassimilate distribution. Indeed, despite unaltered photosynthetic rates, there was a marked shift in carbon partitioning from starch to sugars in the transformants. However, sugars did not greatly accumulate since this was offset by an even greater increase in the loading of assimilates into the phloem. This is an ideal scenario since it circumvents the well characterized phenomenon of feedback inhibition of photosynthesis whereby high leaf sucrose levels suppress the rate of photosynthesis (Stitt, 1990a). In fruits, we demonstrated that the higher sugar influx from leaves led to higher sugar levels in ripe fruit, although significant differences in the main sugars did not correspond to an enhanced Brix index most likely due to corresponding decreases in other components of this composite trait. While enhancing mGS activity into leaf metabolism did not improve CO_2_ assimilation rate, we here demonstrate how metabolic modelling studies can lead to hypotheses generating non-obvious targets for increased crop yield, suggesting that the recently established Crop *in silico* project (https://cropsinsilico.org/) will likely provide considerable novel hypotheses that will ultimately aid us in the metabolic engineering of plants. In our case it suggested that the enhanced mitochondrial activity of glutamate synthetase would represent a route to optimize phloem loading. Our experimental data from plant engineered with this suggested intervention revealed that this was indeed the case and that as a consequence tomato plants expressing either tomato isoform of this enzyme displayed early and increased incidence of flowering, alongside an early fruit set and a shorter life cycle. The fact that at more Northern latitudes the majority of commercial tomato production occurs in greenhouses, moreover, renders the early setting of fruit of potentially highly useful trait. Even more so given that the final fruit yield is additionally enhanced. As such this finding was perhaps surprisingly far more successful than our previous attempts to achieve this using multi-gene approaches (Vallarino et al., 2020).

## METHODS

### Flux balance analysis modelling of tomato leaf metabolism

The model ‘PlantCoreMetabolism v2.0.0’ was based on updated version of our previously described core model of plant metabolism (Shameer et al., 2020) and modified to include gene-to-reaction associations for tomato. A curation log specifying the updates to the model is provided in Supplemental Data Set 4. Gene association data was gathered from PlantCyc Tomato Pathway/Genome Database (www.plantcyc.org). Data from the cropPAL database (Hooper et al., 2016) was used to ensure gene-associations were in-line with known enzyme subcellular localization. The model is provided as an SBML file (Supplemental Data Set 5). A two-phase day-night (diel) formulation was established as previously described (Cheung et al., 2014). The CO_2_ assimilation rate of the model was constrained using the approach previously described (Shameer et al., 2019). The model output was a defined stoichiometry of sugars and amino acids representing tomato phloem sap composition (Walker and Ho, 1977; Valle et al., 1998). Non-growth associated maintenance was set according the incident light flux following a previously described procedure (Töpfer et al., 2020). Other model constraints are listed in Supplemental Table 1. The model was run using the COBRAPy package (Ebrahim et al., 2013) for Python 3. The built-in COBRAPy functions for pFBA and FVA were used to solve the model. A constraint scan on the model output was done by reducing the model output in 2% steps and each time re-running the model with this constrained output and storing the resulting flux solution. Fluxes were graphically plotted using the python package Matplotlib. All code required to run the constraint-scan and generate plots can be found in https://github.com/sshameer/Vallarino_et_al_2021.

### Plasmid construction and generation of transgenic plants

Transformation of tomato plants (*Solanum lycopersicum* cv MoneyMaker) was performed essentially as described in (Nunes-Nesi et al., 2005). Two pK2GW7 plasmids were prepared containing the *nptII* gene for kanamycin resistance, the mitochondrial transit peptide COX4ts (Nelson et al., 2007) and the full-length *S. lycopersicum Glutamine Synthetase (GS)* 1 and 2 genes [*GS1* (*Solyc04g014510*)and *GS2* (*Solyc01g080280*)] in sense orientation under the control of the leaf mesophyll-specific promoter (RbcS). We screened ten and fourteen independent transgenic tomato lines harboring *RbcS-COX4ts-GS1* and *RbcS-COX4ts-GS2* constructs, respectively, based on their levels of *GS1* and *GS2* gene expression. We subsequently selected four lines per construct that were taken to the next generation (T_1_) for detailed characterization. Transgenic plants were grown in greenhouse conditions under 16/8h photoperiod, 60% RH and light intensity 400 µmol m^-2^ s^-1^ as described previously (Nunes-Nesi et al., 2005).

### RNA extraction and gene expression by qRT-PCR

Total RNA was isolated from leaves according to the method described by (Sánchez-Sevilla et al., 2017) Gene expression in tomato leaves was analyzed by real-time quantitative RT-PCR using the fluorescent intercalating dye SYBR Green. Relative quantification of the target genes was performed using the comparative Ct method. Expression data were normalized to the reference genes Ubiquitin3 and elongation factor with comparable results. Values displayed in this publication are normalized to Ubiquitin3 (Wang et al., 2008; Zanor et al., 2009). Primers used are detailed in Supplemental Table 2.

### Photosynthetic parameters

Leaf gas exchange and chlorophyll fluorescence were performed using a Li- 6400XT (Li-Cor Inc., Lincoln, NE, USA). All measurement conditions were as described by (Vallarino et al., 2020).

### Metabolite analysis

The level of starch in 100mg of leaves and 250mg of fruits, respectively, were extracted and analyzed spectrophotometrically as described previously (Dominguez et al., 2013) Pyridine nucleotide levels were performed according to (Schippers et al., 2008). Photosynthetic pigments were determined according to the methods described by Porra et al., (1989). Primary metabolites were carried out in leaves, leaves exudates and pericarp of fruits at different development and ripening stage by using gas chromatography coupled to electron impact ionization-time of flight-mass spectrometry (GC-EI-TOF/MS) according to method described in (Lisec et al., 2006). Absolute metabolite levels in phloem exudate were measured in samples obtained as described in (Lytovchenko et al., 2002), by running standard curves of authentic standards for quantification. Metabolite levels are reported following the reporting standards suggested by (Fernie et al., 2011; Alseekh et al., 2021; Supplemental Data Set 6).

### Mitochondrial isolation

Mitochondria were isolated from 2-month-old tomato leaves as described by Kerbler and Taylor (2017). Briefly, ∼30 g of whole seedlings was disrupted in a precooled mortar and pestle using 150 mL of cold extraction buffer (0.3 M sucrose, 25 mM tetrasodiumpyrophosphate [10 H2O], 2 mM EDTA, 10 mM KH2PO4, 1% polyvinylpyrrolidone [PVP-40], 1% BSA, 20 mM ascorbic acid, 5 mM cysteine [pH 7.5]). The homogenate was filtered twice through four layers of Miracloth (CalBioChem). The material that was retained was recovered and re-ground with 100 mL extraction buffer. This step was repeated three times. The filtrate was first centrifuged at 2,500*g* for 5 min to separate the chloroplast (pellet) and mitochondrial (supernatant) fractions. The supernatant containing the mitochondria-enriched fraction was further centrifuged for 15 min at 18,000*g*. The pellet was resuspended in a small volume of washing buffer (0.3 M sucrose, 10 mM TES-KOH, and 0.1% [w/v] BSA [pH 7.5]). An additional 35 mL of washing buffer was added, and the mixture was centrifuged again at 2,500*g* for 5 min. The supernatant was transferred to new tubes (40 mL), making sure that no pellet was also carried over. The suspension was then centrifuged at 18,000*g* for 15 min. The obtained pellet was resuspended in 2 mL of washing buffer and loaded on top of a 28% (v/v) Percoll (GE Healthcare) continuous gradient with a 0% to 4% (w/v) PVP-40 linear gradient and centrifuged at 38,000*g* for 50 min. The mitochondrial band was collected, washed three times in sucrose wash buffer (without BSA) (0.3 M sucrose, 10 mM TES-KOH [pH 7.5]) and divided into aliquots for measuring enzyme activity.

Purified mitochondria were lysed in the extraction buffer (extraction buffer: 8.7% [v/v] glycerol, 0.5% [w/v] BSA, 1% [v/v] Triton X-100, 50 mM 4-(2-hydroxyethyl)-1-piperazineethanesulfonic acid (HEPES)/KOH pH 7.5, 10 mM MgCl2, 1 mM ethylenediamine tetraacetic acid (EDTA), 1 mM ethylene glycol tetraacetic acid (EGTA), 1 mM benzamidine, 1 mM ɛ-aminocaproic acid, 10 mM phenylmethylsulfonyl fluoride (PMSF) in isopropanol, 0.2 mM leupeptin, and 50 mM dithiothreitol) for mitochondrial enzyme activity measurement (Omena- Garcia et al., 2017).

### Enzyme activities

#### GS

Enzyme activity was measured by the synthetase reaction according to Machado et al., (2001).

#### GOGAT

The GOGAT assay was adapted from (Groat and Vance, 1981) and measured as rate of glutamine-dependent NADH oxidation at 340 nm for 3 min. Extracts were incubated in a solution containing 25 mM KH_2_PO_4_ buffer (pH 7.5), 1 mM hydroxylamine, 2.5 mM 2-oxoglutarate, 0.1 mM NADH, and approximately 100 μg of total protein. The reaction was started by the addition of 10 mM of glutamine.

#### GDH

GDH was measured as described by (Robinson et al., 1991), as the rate of 2- oxoglutarate-dependent NADH oxidation at 340 nm for 3 min.

#### PGM, PEPC, NAD-MDH and NADP-MDH

Enzyme extracts were prepared as described previously by (Gibon et al., 2004).

#### AGPase

Enzyme activity was assayed as described in (Osorio et al., 2013).

#### NR and NiR

NR was determined as decreased absorbance at 340 nm due to NADH oxidation as described in (Campbell and Smarrelli, 1978). NiR activity was measured exactly as described previously (Lillo and Ruoff, 1992). The absorbance was read at 540 nm at 25°C.

#### SDH and UGPase

Total protein extracted from purified mitochondria were used to analyze SDH and UGPase described previously (Omena-Garcia et al., 2017).

### Analysis of the redistribution of positionally labelled Glc

TCA cycle flux analysis was calculated based on ^14^CO_2_ evolution. Fruit pericarp disks at 22 DAP stage from transgenic lines and WT (10 disks per biological replicate. A total of 5 replicates were used) were incubated in 5 ml of 10 mM MES-KOH, pH 6.5 containing 2.85 kBq ml^-1^ of [1-^14^C]- or [3,4-^14^C] Glc. Evolved ^14^CO_2_ was collected in 0.5 ml KOH (10% w/v) every hour (for 5 h) and quantify by liquid scintillation counting. Disks were washed five times in 10 mM MES- KOH, pH 6.5 and frozen in N_2_ liquid for further analysis. The ^14^C-labeled material was fractionated according to (Lytovchenko et al., 2002). The results were interpreted following recommendations detailed by (Geigenberger et al., 2000).

### Agrobacterium culture preparation and infiltration for mitochondrial localization

The agrobacteria were grown on YEB-induced medium plates (Zhang et al., 2020) (at 28°C for 24-36h. The cells were scratched and resuspended in 500µl washing solution (10mM MgCl_2_, 100μM acetosyringone). After briefly vortexing, the 100µl resuspended agrobacteria were diluted 10 times to measure the OD600 (should be equal to or greater than 12). The agrobacteria were finally diluted to OD600 = 0.5 in infiltration solution (¼MS [pH=6.0], 1% sucrose, 100μM acetosyringone, 0.005% (v/v, 50μL/L) Silwet L-77). The agrobacteria were infiltrated into tobacco leaves kept in the light to dry the leaves (1 hour), using a 1 ml plastic syringe, and then subsequently kept in the dark for 24 hours at room temperature. The transformed plants were then transferred back to the greenhouse for another 2-3 days before microscopy.

## Accession numbers

Sequence data for GS1 and GS2 genes can be found in the solcyc.solgenomics database under the following numbers: *GS1* (*Solyc04g014510*) and *GS2* (*Solyc01g080280*).

## Supplemental Data

**Supplemental Figure 1.** Predicted glutamine synthase fluxes in mature tomato leaves as the export rate of sugar and amino acids to the phloem was varied between 60 – 100% of the maximum achievable within the model constraints.

**Supplemental Figure 2.** Flower and ripe fruit apperance in RbcS–GS1 and RbcS–GS2 lines.

**Supplemental Figure 3.** Phenotypic characterization of the *RbcS–GS1* and *RbcS–GS2* lines grown in Autumn season.

**Supplemental Figure 4.** Photosynthetic parameters in *RbcS–GS1* and *RbcS– GS2* lines.

**Supplemental Figure 5.** Expression of tomato *GDC-L gene* by qRT-PCR.

**Supplemental Table 1.**List of constraints applied to the diel stoichiometric model of tomato leaf metabolism.

**Supplemental Table 2**. List of oligonucleotides used in this study.

**Supplemental Data Set 1.** Full dataset of the flux distributions.

**Supplemental Data Set 2**. Metabolite contents in leaves from RbcS–COX4ts- GS1 and RbcS–COX4ts-GS2 lines.

**Supplemental Data Set 3**. Absolute metabolite concentrations (nmol/g DW) of diurnal period exudates from leaves of five-week old transgenic plants and WT were collected for 8h.

**Supplemental Data Set 4.** List of changes to PlantCoreMetabolism since previously published version 1.2.3.

**Supplemental Data Set 5**. Model from flux balance analysis provided as an SBML file.

**Supplemental Data Set 6**. Metabolite Reporting Guidelines.

## Acknowledgements

We acknowledge the excellent care of the plants by GreenTeam (Max Planck Institut für Molekulare Pflanzenphysiologie). The work of the authors is supported by funding from the Max-Planck Society (JGV, Y.Z., and A.R.F). A.R.F and Y. Z would like to thank the European Union’s Horizon 2020 research and innovation programme, project PlantaSYST (SGA-CSA No. 739582 under FPA No. 664620) for supporting their research.

## Author contributions

J.G.V, S.S, L.J.S. and A.R.F designed the research. S.S, R.G.R, and L.J.S. performed the flux balance analysis. J.G.V generated the constructs, generated transgenic plants, performed and analyzed most experiments and data. Y.Z. and J.G.V. performed confocal microscopy experiment. J.G.V and A.R.F wrote the article. All authors commented on the article.

